# Male mate preference against tail-elongated females in the barn swallow

**DOI:** 10.1101/2022.05.22.492952

**Authors:** Masaru Hasegawa, Emi Arai

## Abstract

The function of female ornamentation in the context of (inter)sexual selection attracts keen attention these days, but empirical field studies are mostly based on indirect measures such as mating patterns; direct evidence of male mate preference on female ornamentation while controlling for confounding factors is needed. Here we performed model presentation experiments to study male mate preference in the barn swallow *Hirundo rustica*, a model species of sexual selection. Female barn swallows have somewhat shorter tails than males, but their tails are still long and costly. Although many correlational and experimental studies of live females have focused on long tails in female barn swallows over the last three decades, direct behavioral tests of male mate preference are lacking. By using a sequential model presentation experiment, in which we repeated the trials with the same males, we found that males significantly reduced the number of pairing displays when they were presented with tail-elongated female models compared to control female models. Interaction between treatment and male tail length was far from significant. The observed pattern is inconsistent with the prevailing view that costly female ornamentation is maintained via male preference for more ornamented females. Rather, the observed pattern is consistent with the sexual mimicry hypothesis, in which females can avoid sexual (and social) harassment by mimicking males.

## 1 INTRODUCTION

Animals often exhibit ornamentation, such as colorful bird plumage and long tail feathers, which seems to have rather negative effects on survivorship (Andersson 1994). A classic explanation of such ornamentation is that more ornamented individuals are preferred by potential mates and hence sexually selected (e.g., see Hill 2006 for reviews on the plumage characteristics of birds). Although this explanation is often applied to male ornamentation, not only males but also females possess ornamentation. Recent phylogenetic and experimental studies have revealed that the correlated response hypothesis, i.e., intersexual genetic correlation with male ornamentation, alone cannot explain female ornamentation (e.g., Dale et al. 2015), and that female ornamentation would evolve and be maintained due to its function in social interactions (reviewed in Amundsen & Pärn 2006; Kraaijeveld et al. 2007; Tobias et al. 2012; Hare & Simmons 2019; Hernandez et al. 2021).

The function of female ornamentation, however, remains unclear, because most field studies report indirect measures of male mate preference, such as breeding onset date, assortative mating, and so on (reviewed in Amundsen & Pärn 2006). These indirect measures are not a strong test of male mate preference, because male mate preference is at best of minor importance in determining these measures (compared to other factors, such as female age, physiological condition, arrival date, resource availability, and so on: e.g., see Amundsen 2000; Amundsen & Pärn 2006; Kraaijeveld et al. 2007 for reviews). This is also the case even when female ornamentation is experimentally manipulated, because ornamentation feeds back to physiological state (e.g., Vitousek et al. 2013) and behavior (e.g., ornamentation and its manipulation may affect general and mating activity; e.g., Barnald 1990; Balmford & Thomas 1992; Wolf et al. 2004), which confounds indirect (and even direct) measures of male mate preference. Male mate preference on female ornamentation measured by male behavior while controlling for these confounding factors, for example, using model presentation experiments, provides direct evidence of male mate preference, though only a handful studies are reported (e.g., Jones & Hunter 1999; reviewed in Amundsen & Pärn 2006; Hernandez et al. 2021).

The barn swallow *Hirundo rustica* is a socially monogamous bird and a model species of sexual selection, in which females have been shown to behaviorally prefer tail-elongated males, explaining their long outermost tail feathers (e.g., acceptance/rejection of extra-pair copulation; Møller 1988; reviewed in Møller 1994; Turner 2006; Romano et al. 2017; also see Hasegawa & Arai 2020a for comparative support for the evolution of tail fork depth in relation to high opportunities of extrapair paternity in hirundines). Compared to males, female barn swallows have somewhat shorter tails, but their tails are nonetheless costly to produce and maintain as is the case for male tails (e.g., Cuervo et al. 2006; Cuervo & de Ayala 2014; Hasegawa et al. 2020; also see Hasegawa & Arai 2017 for comparative support); thus, a selective advantage is needed to explain their existence (Amundsen 2000). In fact, in hirundines (family: Hirundinidae), male and female tails evolved differently (Hasegawa & Arai 2020b) and hence sexual dimorphism in tail length varies greatly across species (Turner & Rose 1994; Hasegawa & Arai 2020a), indicating that the correlated response hypothesis alone cannot explain the expression of female tail length. Many correlational studies support the sexual selection explanation for the evolution of long female tails (e.g., Møller 1993a, 1994; see Hasegawa et al. 2017 for Japanese barn swallows), but, in contrast to these correlational studies, manipulative experiments on female tails do not support sexual selection for long tail feathers (Cuervo et al. 1996; also see Hasegawa et a. 2018, 2020 for manipulative experiments showing no detectable differential allocation). However, all these correlational and experimental studies used indirect measures of mate preferences (i.e., breeding onset date, assortative mating, paternal investment, annual reproductive success; reviewed in Romano et al. 2017), and thus cannot be conclusive (sensu Amundsen & Pärn 2006; see above). In highly aerial birds like barn swallows, indirect measures of mate choice should also be affected by the aerodynamic cost of long female tails (e.g., Buchanan & Evans 2000; Hasegawa et al. 2020), preventing us from elucidating the function of long tails in females. In fact, “any experimental shortening and lengthening of the outer tail feathers is likely to upset an original co-adapted character set…” (Norberg 1994, p. 231), which would then affect female physiology and behavior, possibly confound with indirect measures of mate choice. To date, no studies reported direct male mate preference in relation to female tail length in this model species of sexual selection (see Romano et al. 2017 for all sexual selection studies in this species up to the publication year).

Here, using model presentation experiments (Hasegawa et al. 2016a), we tested male mate preference for a plumage ornament, a long tail, in the barn swallow in Japan (subspecies: *gutturalis*). Because unpaired male barn swallows lead females to their nest sites with characteristic enticement calls (hereafter “pairing display”; Hasegawa et al. 2013; reviewed in Møller 1994; Turner 2006; Hasegawa 2018; also see supplementary videos), male mate preference for female models can be evaluated easily based on the number of pairing displays (Hasegawa et al. 2016a). We presented control or tail-elongated stuffed female models toward naïve males (i.e., single-female experiment; Houde 1997). A few days after the first trial, we performed a second trial using alternative models (i.e., tail-elongated models to males previously exposed to controls and vice versa). By using this “sequential” model presentation procedure, we can examine individual mate preferences of males regarding female tail length. We conducted this “sequential choice” test rather than the “simultaneous choice” test, because the former is more appropriate than the latter in the barn swallow, a protandry migrant species, in which males encounter females sequentially rather than simultaneously as in many other species (Barry & Kokko 2010). Because long-tailed females are high-quality females in terms of arrival date, survivorship, breeding experience, and parental and reproductive investment in barn swallows including Japanese populations (e.g., Møller 1993a, 1994; Brown & Brown 1999; Hasegawa et al. 2016b, 2017, 2018, 2020), and because previous correlational studies support sexual selection for female long tails in Japanese barn swallows (Hasegawa et al. 2017), we predicted that males would prefer tail-elongated female models to control female models. We also examined individual mate preference for female tail length in relation to male tail length, which is virtually impossible to examine when using indirect measures of mate preference (e.g., breeding date, assortative mating; see above). Sexual selection theory predicts a positive correlation between male tail length and male mate preference for female tails (e.g., prudent mate preference: Härdling & Kokko 2005; reviewed in Pollo et al. 2022). We discuss potential explanations for the patterns observed in the present study and the function of female ornamentation.

## 2 METHODS

### 2.1 Study site

This study was conducted in 2015 in a residential area of Hayama-machi, Kanagawa Prefecture (35°16′N, 139°35′E) and in 2020–2022 in a residential area of Tsurugi-machi, Ishikawa Prefecture (36°26′N, 136°37′E), Japan. In these areas, swallows nest sparsely under the eaves of a covered sidewalk along the street (Tajima & Nakamura 2003; Hasegawa & Arai 2016), which enabled us to quantify individual male responses to each model without confounded by neighboring males. In these sparsely breeding barn swallows, extrapair paternity is virtually absent (<3%: Hasegawa et al. 2010a). We focused on unpaired males soon after their arrival at the study site through daily observation (see Arai et al. 2009 for detailed methodology). Male identity was confirmed based on a unique combination of two or three colored rings that were attached previously (e.g., see Fig. S1 for color-ring based identification), or by the position of breeding territory and male morphology (e.g., very long tails, notable tail asymmetry, large throat patches, and so on; note that territory takeover is rare in Japanese populations; Hasegawa & Arai 2016). In the latter cases, all males were captured and ringed after they initiated reproduction in the same territory (see Hasegawa et al. 2010b for detailed methodology). When we captured and ringed males, we measured male tail length and other morphological traits (though we here focused on male tail length alone; see Hasegawa et al. 2010b for detailed information).

### 2.2 Model presentation

As before (Hasegawa et al. 2016a), we used stuffed female swallows (i.e., models) to study male mate preference. To minimize pseudoreplication, we used five different models, which were made from females that were found to be dead or accidentally died during previous study years. For tail-elongated treatment, we elongated the outermost tail feathers by ca. 17 mm (range: 16–19 mm when measured), which is roughly comparable to the elongation in a previous manipulative experiment (20 mm elongation compared with controls; Cuervo et al. 1996), by attaching additional tail feathers plucked from other females with thin glue tapes (Fig. 1; also see Fig. S1). As in the Spanish population of Cuervo et al. (1996), female tails in Japanese barn swallows are ca. 15% shorter than male tails, but still exhibit high variance (e.g., during the 2005 breeding season in the Joetsu population, in which the largest samples could be obtained by intensive field survey, males: mean ± SD = 94.42 ± 8.16, range = 75.13, 119.12, *n* = 164; females: 79.93 ± 4.58, range = 66.76, 96.89, *n* = 179; see Hasegawa et al. 2010b for detailed methodology; note that we were able to capture only a small portion of the whole populations in the current study, which prevents accurate estimation of the distribution of tail length at these sites, although we observed similar mean ± SD in these populations; e.g., Hasegawa et al. 2016b). Because the tail lengths of the five stuffed females were ca. 71 mm, 73 mm, 74 mm, 76 mm, and 82 mm, the tail-elongated treatment roughly match the range of natural tail lengths in Japanese barn swallows (note that the accuracy of measurements of stuffed females was limited compared with measurements of live females; Hasegawa & Arai 2013). To reduce pseudoreplication, we used tail feathers plucked from two different females and randomly used one of the two sets of tail feathers. For the control treatment, we used stuffed models without additional tail feathers (Fig. 1).

**FIGURE 1.**
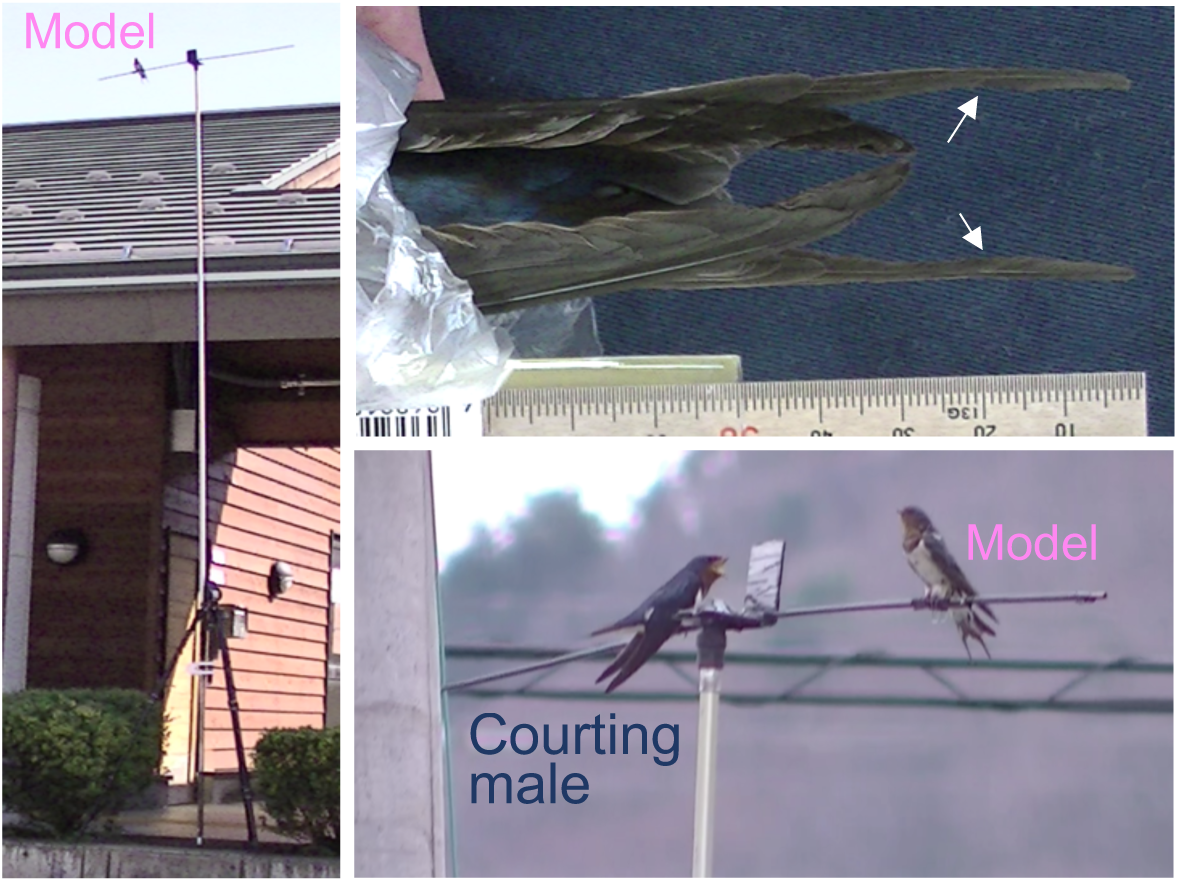
Experimental setup of model presentation in the current study. Left panel shows the entire setup. Model was presented ca. 5 m above ground. Upper right panel shows tail-elongated female model (arrows indicate the end of the original outermost tail feathers). Lower right panel shows a focal male displaying a typical courtship posture (left) toward a control female model (right, picture captured from a videotape, which is included as a supplementary material; note that the male is lowering its wings below the tail feathers)

We conducted experiments under sunny or cloudy weather conditions (i.e., when we could observe male behavior adequately) in March to April. In the first trial, we randomly presented each unmated male with a control or tail-elongated model. We presented the model on a horizontal wire 5 m above the ground in the male’s territory (Fig. 1; see Hasegawa et al. 2016a for a schematic illustration). During 10-min presentation of the model to the focal male, we recorded the number of copulation attempts and the number of pairing displays (i.e., nest visiting, often with enticement calls; see fig. 1 in Hasegawa 2018 for courtship sequence; Hasegawa et al. 2013; see supplementary videos). Because males rarely attempted copulation in the current study, we focused on pairing displays. In fact, pairing displays, but not copulation attempts, were related to a female trait (ventral plumage coloration) in the preceding study, perhaps because males indiscriminately copulate with females (Hasegawa et al. 2016a). Nevertheless, information on copulation attempts would be helpful, and thus we presented the data together with the analyses (note that all analyses were far from significant; see results). Because pairing displays are not associated with copulation attempts (Møller 1994; Turner 2006; Hasegawa 2018), these two measures are largely independent (Hasegawa et al. 2016a).

We performed the second trial a few days (ca. three days; range: 1–9 days) after the first trial to study whether the behaviors of each male changed with treatment. This was possible only when males remained unmated under favorable weather conditions (note that mating status was surveyed every day; see above). Alternative models were presented to each male in the same location during the second trial (i.e. tail-elongated models to males previously exposed to control models and vice versa). As in the first trial, we recorded the number of copulation attempts and pairing displays over 10 min. To detect subtle male mate preference, we used the same female model in the first and the second trial, which was shown to be effective in barn swallows (i.e., high detectability of male mate preference: Hasegawa & Arai 2016).

### 2.3 Statistics

A Bayesian generalized linear mixed model (MCMCglmm function in the R package MCMCglmm) with a Poisson error distribution was used to test the effects of treatment (“control” and “tail-elongated” are denoted as 0 and 1, respectively) on the number of male pairing displays (prior: [list(G = list(G1 = list(V = 1, nu = 0.002)), R = list(V = 1, nu = 0.002)). We did not include the natural tail length of the stuffed models as a covariate due to the small variation in female tail length in the models, i.e., ca. 10 mm difference between shortest and longest tail, which is one-third of the natural variation in tail length in Japanese barn swallows (see the preceding section). It should be noted that the MCMCglmm function accounts for overdispersion by using a random factor, “units” (Hadfield 2010). We included male identity as a random factor to study changes in behavior (note that we did not test the same males across years). We did not include the identity of stuffed females as another random factor in these models not to complicate the model structure (although we confirmed that including this variable did not change the results qualitatively; see results). Based on the previous model presentation experiment in Japanese barn swallows (see table 2 in Hasegawa et al. 2016), repeated model presentation using 22 different males provided adequate statistical powers (89.6%, based on “powerSim” function while controlling for overdispersion using observation level random effect in the package “simr”: Green & MacLead 2016), though actual statistical power would be lower, because morphological differences between treatments were much smaller in the current study than in the previous one. This is in fact the reason why we focused solely on repeated presentation as a sequential mate preference (rather than on a single, first-time presentation). We ran for 520,000 iterations, with a burn-in period of 120,000 and a thinning interval of 200 in each iteration. The reproducibility of the MCMC simulation was assessed by calculating the Brooks–Gelman–Rubin statistic (Rhat), which must be <1.2 for all parameters (Kass et al. 1998), by repeating the analyses three times. All data analyses were performed using the R (version 4.0.3) statistical package (R Core Team 2020).

### 3 RESULTS

The behaviors of 22 males (*n* = 1 in 2015, *n* = 5 in 2020, *n* = 7 in 2021, and *n* = 9 in 2022) were repeatedly observed in two trials conducted on different dates (and thus the total number of trials was 44). The number of pairing displays was significantly explained by treatment (Table 1 above): males decreased the number of pairing displays when presented with a tail-elongated female model (Fig. 2). The number of copulation attempts did not significantly change with treatment (Table 1 below; Fig. 2). Presentation order was not significant in either model (Table 1). Including the identity of female model as another random factor did not change the results qualitatively (i.e., significant and non-significant variables remained unchanged; data not shown).

**TABLE 1.** Bayesian generalized linear mixed model with a Poisson error distribution explaining (a) the number of male pairing displays and (b) the number of copulation attempts during repeated trials in male barn swallows (*n*_*male*_ = 22, *n*_*total*_ = 44)

| Variable | Coefficient [95% CI] | $pMCMC$ |
| --- | --- | --- |
| (a) pairing displays |  |  |
| Intercept [control] | 1.51 [1.01, 1.93] | **<0.0001** |
| Treatment [elongated – control] | -0.58 [-0.89, -0.26] | **0.001** |
| Order of presentation [2<sup>nd</sup> – 1<sup>st</sup>] | -0.23 [-0.54, 0.10] | 0.15 |
| (b) copulation attempts |  |  |
| Intercept [control] | -2.07 [-4.47, -0.19] | **0.01** |
| Treatment [elongated – control] | -0.41 [-1.34, 0.54] | 0.35 |
| Order of presentation [2<sup>nd</sup> – 1<sup>st</sup>] | 0.03 [-0.82, 0.96] | 0.96 |

We included male identity (variance = 0.82 and 8.25) as a random factor in addition to “units,” which deal with overdispersion (variance = 0.03 and 0.33) for the analyses of the number of pairing displays and copulation attempts, respectively. Estimated repeatabilities after controlling for covariates were 0.75 (95% CI = 0.53, 0.90) and 0.71 (95% CI = 0.40, 0.91) for the number of pairing displays and copulation attempts, respectively (see Nakagawa & Schielzeth 2010 for calculation).
Note: Mean coefficients, 95% credible intervals (CI), and MCMC-based $p$ -values are shown. Significant test results (i.e., $pMCMC < 0.05$ ) are indicated in bold

**FIGURE 2.**
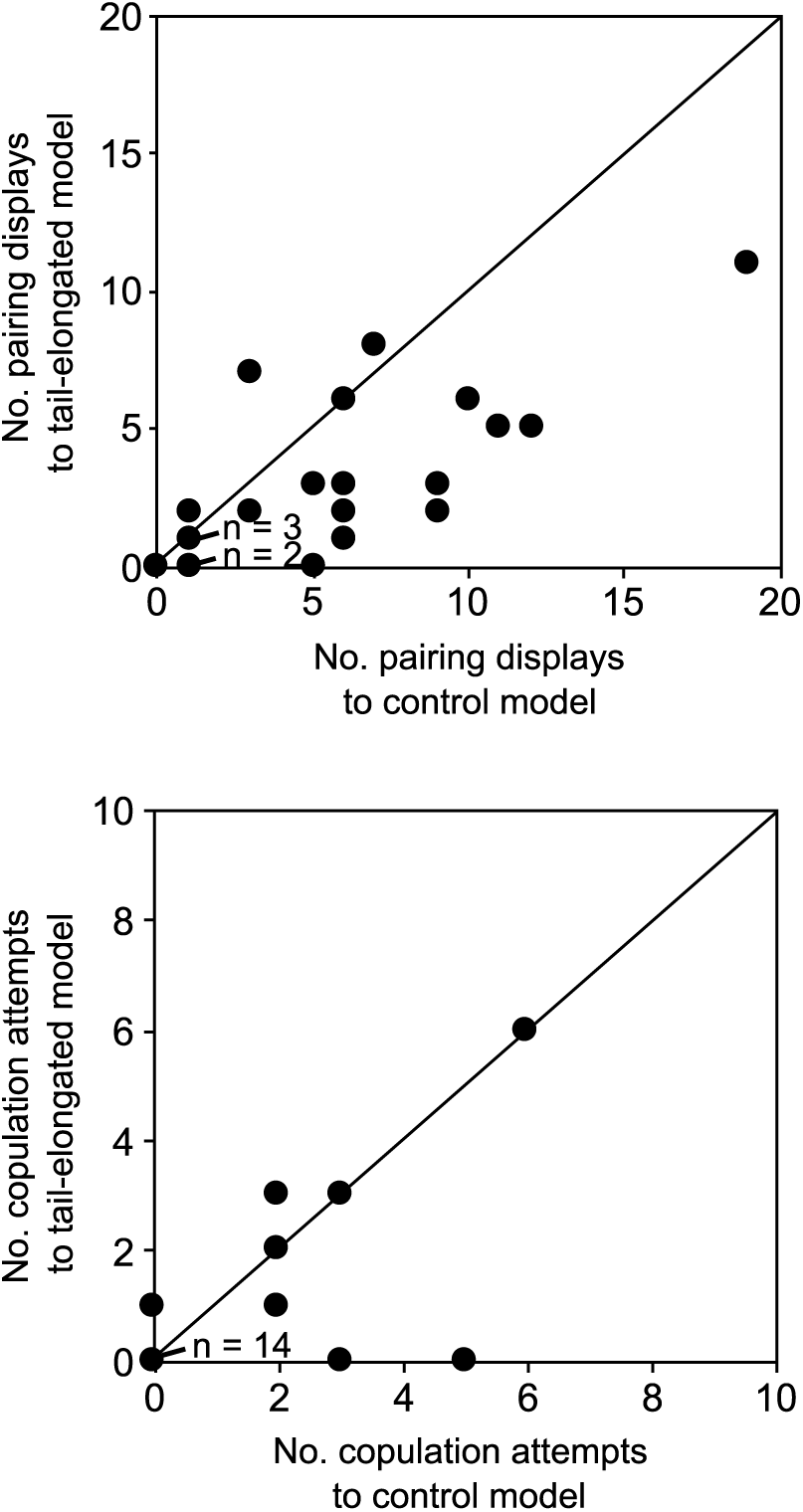
Relationship between the number of male responses when presenting the control female model (x-axis) and when presenting the tail-elongated female model (y-axis) in barn swallows (upper panel: male pairing displays; lower panel: male copulation attempts). The diagonal line indicates identical values

Using a subset of data, in which male morphological measurements were available (*n*_*male*_ = 16), we tested the interaction between male tail length and treatment. Male tail length did not significantly explain the number of pairing displays with and without including the interaction term between treatment and male tail length (Table S1). Similar non-significant patterns were found when we analyzed the number of copulation attempts (Table S1).

## 4 DISCUSSION

The main finding of the current study is that, contrary to our prediction, male barn swallows did not show a preference for tail-elongated female models. Rather, our model presentations showed male mate preference *against* tail-elongated female models, irrespective of their own tail length. To our knowledge, this is the first direct test of male mate preference on female tail length in this model species of sexual selection. This pattern sharply contrasts with correlational studies, which suggest sexual selection for long-tailed females (e.g., Møller 1993a; also see Hasegawa et al. 2017 for Japanese barn swallows), and with manipulative experiments on live females, which found no detectable effects of treatment on reproductive performance, including breeding onset date (Cuervo et al. 1996). Given that indirect measures of mate choice (e.g., breeding onset date) are affected by several confounding factors (see Introduction), the apparent inconsistency between previous studies and the current behavioral test is not surprising. Still, male mate preference against well-ornamented females, which has been reported in only a few cases in birds (e.g., bill color preferences of zebra finches; Burley & Coopersmith 1987), needs explanation.

First of all, it should be noted that our prediction that males should prefer long-tailed females was based on the assumption that male barn swallows can easily distinguish females from males. Even if long-tailed females are favorable mates, males might avoid male-like females and prefer typical females (i.e., females with average-sized ornaments) not to waste time and energy by courting on (and then mate-guarding of) potential rivals. This possibility is likely, because long-tailed females resemble males more than short-tailed females (Fig. S2). Although male and female barn swallows differ in several aspects, including dorsal and ventral plumage coloration, tail length is the most notable difference between sexes (Turner 2006; note that weak sexual dimorphism does not function in sex identification; Burley 1981). Another model presentation experiment, in which stuffed male models were used, showed that male barn swallows even copulate with male models (Brodetzki et al. 2021), supporting the difficulty for males in distinguishing females from males.

The sexual mimicry hypothesis predicts that females produce male-like characteristics to avoid costly male sexual harassment (reviewed in Amundsen 2000; Amundsen & Pärn 2006; Kraaijeveld et al. 2007; Tobias et al. 2012). Although vertebrate researchers often assume fitness benefits of female attractiveness (but see Burley 1981), avoiding male attention (i.e., being unattractive) is a well-known function of male-like characteristics in invertebrates (e.g., Robertson 1985; see reviews listed above; also see Falk et al. 2021 suggesting this function in birds). As we showed here, tail-elongated female models elicited male responses less than control models but still attracted the attention of males, and thus the observed unattractiveness would not be costly for females. Rather, even in just 10-min observations (see methods), we observed ca. four-time male pairing courtship displays toward female models (Fig. 2; also see supplementary videos for actual male behavior), and thus sexual mimicry of males, which can persist throughout the breeding season unless individuals reveal their sexes behaviorally, could be adaptive by somewhat reducing male harassment. Sexual conflict over courtship interactions, rather than sexual selection, might explain female ornamentation, particularly in this relatively high-density species that encounter many conspecifics while foraging (note that this situation can favor the evolution of sexual mimicry; Burley 1981). Such benefits of ornamentation, perhaps together with intersexual genetic correlation, explain female ornamentation (note that, because intersexual genetic correlation itself can evolve, this factor alone cannot explain costly female ornamentation: Amundsen & Pärn 2006).

An alternative explanation is that, even if males somehow distinguish females from males, males might avoid long-tailed females for other reasons. For example, female ornamentation might trade off with maternal investment and thus males might avoid highly ornamented females to ensure their own reproductive success (Chenoweth et al. 2006). This explanation seems unlikely given that naturally long-tailed females show higher (rather than lower) reproductive investment (see Introduction). However, egg size decreases with female tail length at least at interspecific levels in hirundines (Hasegawa & Arai 2017) and the length of the incubation period increases with female tail length in Japanese barn swallows (Hasegawa et al. 2018), both of which cannot easily be compensated by their mates (note that male barn swallows lack brood patches; Turner 2006; Voss et al. 2008). Thus, male mate preference against long-tailed females might be beneficial to avoid inevitable loss of reproductive (maternal) investment in advance, though the importance of this kind of reproductive benefit might be relatively small compared to the benefit of pairing with long-tailed females (see above).

In addition, in Asian barn swallows (*H. r. gutturalis*), which have shorter tails than European barn swallows (*H. r. rustica*; Turner 2006), mate preference against long-tailed females might be adaptive to avoid interbreeding (Servedio 2007), particularly in the barn swallow in which females disperse more than males (and thus males may encounter immigrant females, possibly across continents: Zink et al. 2006; Hasegawa et al. 2016a). Even within Asian subspecies, Japanese barn swallows have shorter tails than continental ones (e.g., Chinese barn swallows; Liu et al. 2018; M Hasegawa unpublished data), and thus local adaptation would favor male xenophobia (i.e., male avoidance of long-tailed females). However, the importance of this factor would be limited, because the proportion of such long-distance migrants from other regions would be negligible in the population. No detectable relationship between treatment and male tail length further indicates that a simple assortative mate preference is unlikely. Finally, in addition to these “adaptive” male mate choices, non-adaptive mate choice can persist in the population, for example, due to genetic correlation to other adaptive traits (Amundsen & Pärn 2006), which should be tested in future. It should be noted that these alternative explanations for male mate preferences against long-tailed females can explain the observed male behavior, but, unlike the sexual mimicry hypothesis, are insufficient to explain why females possess long tails. The current study, however, focused solely on male mate preference and thus long, costly tails might be selected for in other contexts, such as female-female competition for mates and non-sexual social selection (e.g., as written in Burley 1981, females might bluff about their territory defense ability by possessing male plumage; also see Falk et al. 2021 for the function of female ornaments to reduce social harassment; reviewed in Tobias et al. 2012).

A caveat of the current study is that male barn swallows might distinguish experimentally elongated tails from naturally long tails. We could not completely exclude this possibility. However, this is unlikely, because tail-elongated models look quite similar to naturally long-tailed females, at least to human observers (Fig. 1 & S1; also see supplementary videos), and much more casual manipulation is reported to be effective, at least when manipulating male tail feathers (e.g., cut-and-glue, white painting, and so on; e.g., Møller 1988, 1993b). Or, because any behavioral components of females were excluded in our model presentation, this might have affected the outcome of the experiments. This possibility is also unlikely because females normally perform no courtship displays during male pairing attempts (reviewed in Hasegawa 2018).

In conclusion, we observed male mate preference against well-ornamented females in the barn swallow, a model species of sexual selection. In addition to adaptive and non-adaptive mate preference, the sexual mimicry hypothesis, which is often ignored but can explain female ornamentation in the context of sexual conflict, should be carefully argued in future. As shown here, model presentation experiments can capture subtle male mate preference and thus represent a useful methodology in studies of female ornamentation. Such an experiment would be particularly valuable when testing individual male mate preferences in relation to their own phenotype (ornamentation, here), which has still been rarely performed in the wild (Pollo et al. 2022) and thus general pattern remains to be clarified.

## ACKNOWLEDGMENTS

We are grateful to the residents of Hayama-machi and Tsurugi-machi for their kind support and assistance. We thank Dr. Nobuyuki Kutsukake and his laboratory members at the Sokendai (Graduate University for Advanced Studies) for their kindest support. We also thank Dr. Shumpei Kitamura and his laboratory members in Ishikawa Prefectural University. MH was supported by the Research Fellowship of the Japan Society for the Promotion of Science (JSPS, 15J10000; 19K06850, 22J40066).

## AUTHOR CONTRIBUTION

MH did most field survey, performed statistical analysis, and wrote the most of the manuscript, EA assisted field survey and improved the manuscript.

## ETHICAL APPROVAL

The permits for the current study including capturing were provided by Kanagawa Prefecture in Japan (#26-0130-01) and Sokendai (2014A005) and by Ishikawa Prefecture in Japan (#19109, #20113, #20249) and Ishikawa Prefectural University (#R2-14-1, #R3-14-3, #R4-14-2), following the Wildlife Protection and Hunting Management Law.

## DATA AVAILABILITY STATEMENT

Data will be deposited into osf.io once accepted (we also included the data in Table S2).

---

**TABLE S1.** Bayesian generalized linear mixed model with a Poisson error distribution explaining (a) the number of male pairing displays and (b) the number of copulation attempts in relation to tail length and its interaction with treatment during repeated trials in male barn swallows (*n*_*male*_ = 16, *n*_*total*_ = 32)

| Variable | With interaction |  | Without interaction |  |
| --- | --- | --- | --- | --- |
|  | Coefficient [95% CI] | *p*MCMC | Coefficient [95% CI] | *p*MCMC |
| (a) pairing displays |  |  |  |  |
| Intercept [control] | **1.24 [0.64, 1.89]** | **<0.01** | **1.21 [0.55, 1.87]** | **<0.01** |
| Treatment [T: elongated – control] | **-0.83 [-1.30, -0.34]** | **<0.01** | **-0.76 [-1.17, -0.30]** | **0.001** |
| Order of presentation [2<sup>nd</sup> – 1<sup>st</sup>] | 0.01 [-0.46, 0.43] | 0.94 | 0.00 [-0.40, 0.42] | 0.97 |
| Male Tail length [M] | 0.31 [-0.33, 0.96] | 0.30 | 0.35 [-0.30, 0.93] | 0.25 |
| Interaction [TxM] | 0.12 [-0.34, 0.54] | 0.55 | — | — |
| (b) copulation attempts |  |  |  |  |
| Intercept [control] | **-3.41 [-8.07, 0.09]** | **0.02** | **-3.70 [-10.04, 0.12]** | **0.02** |
| Treatment [T: elongated – control] | -0.97 [-3.8, 1.19] | 0.25 | -0.76 [-2.89, 1.59] | 0.34 |
| Order of presentation [2<sup>nd</sup> – 1<sup>st</sup>] | 0.10 [-2.13, 2.46] | 0.98 | 0.31 [-1.64, 3.09] | 0.81 |
| Male Tail length [M] | 0.44 [-2.67, 4.00] | 0.71 | -0.02 [-3.66, 3.91] | 0.95 |
| Interaction [TxM] | -1.65 [-5.23, 1.23] | 0.18 | — | — |

We included male identity (variance = 1.14 and 20.25) as a random factor in addition to “units,” which deal with overdispersion (variance = 0.06 and 4.55) for the analyses of the number of pairing displays and copulation attempts when including interaction terms, respectively. When excluding interaction terms, variance of male identity and units were as follows (male identity: 1.19 and 24.75; units: 0.05 and 5.14), respectively.
Note: Mean coefficients, 95% credible intervals (CI), and MCMC-based $p$ -values are shown. Significant test results (i.e., $p_{MCMC} <$ 0.05) are indicated in bold

**FIGURE S1.**
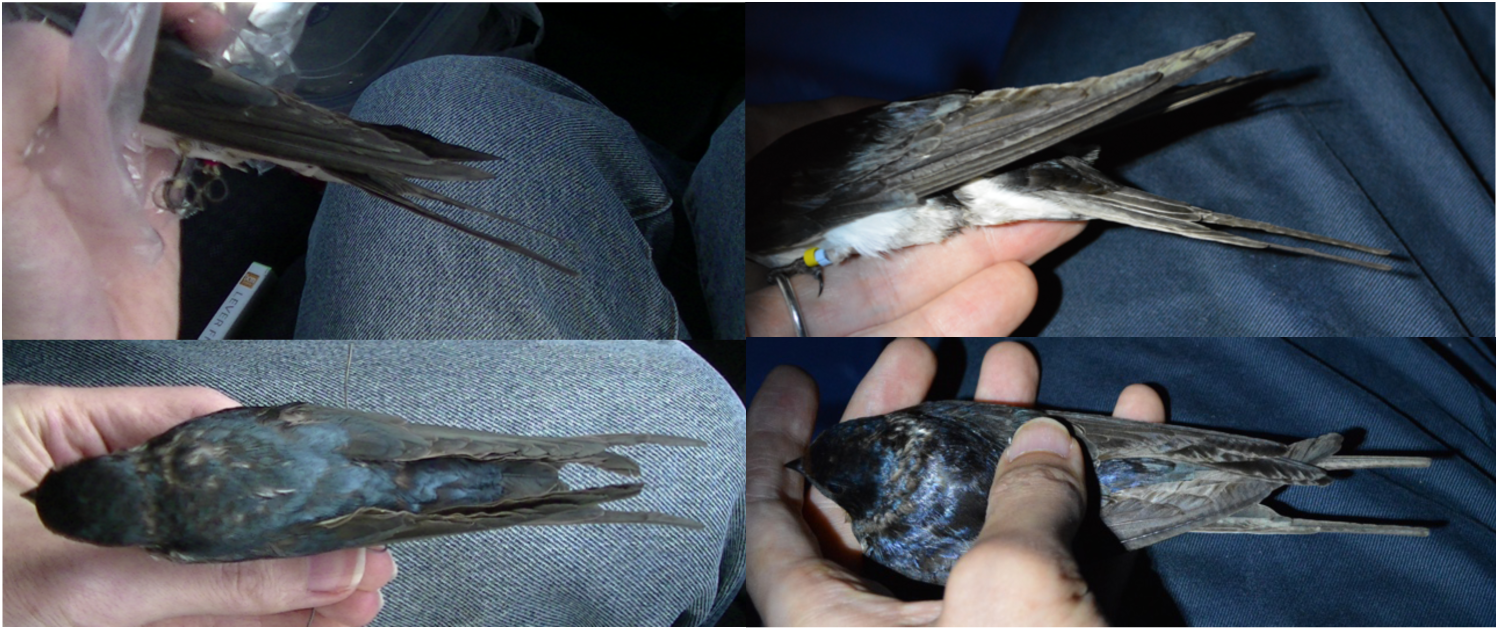
Photographs of a tail-elongated female model (left) and a naturally long-tailed female of the Tsurugi population (right). The photographs show a close-up view from the side (above) and a whole-body picture (below). Note that different female models are shown in the left pictures and that long-tailed female in the right picture had somewhat fatty body shape in the right picture)

**FIGURE S2.**
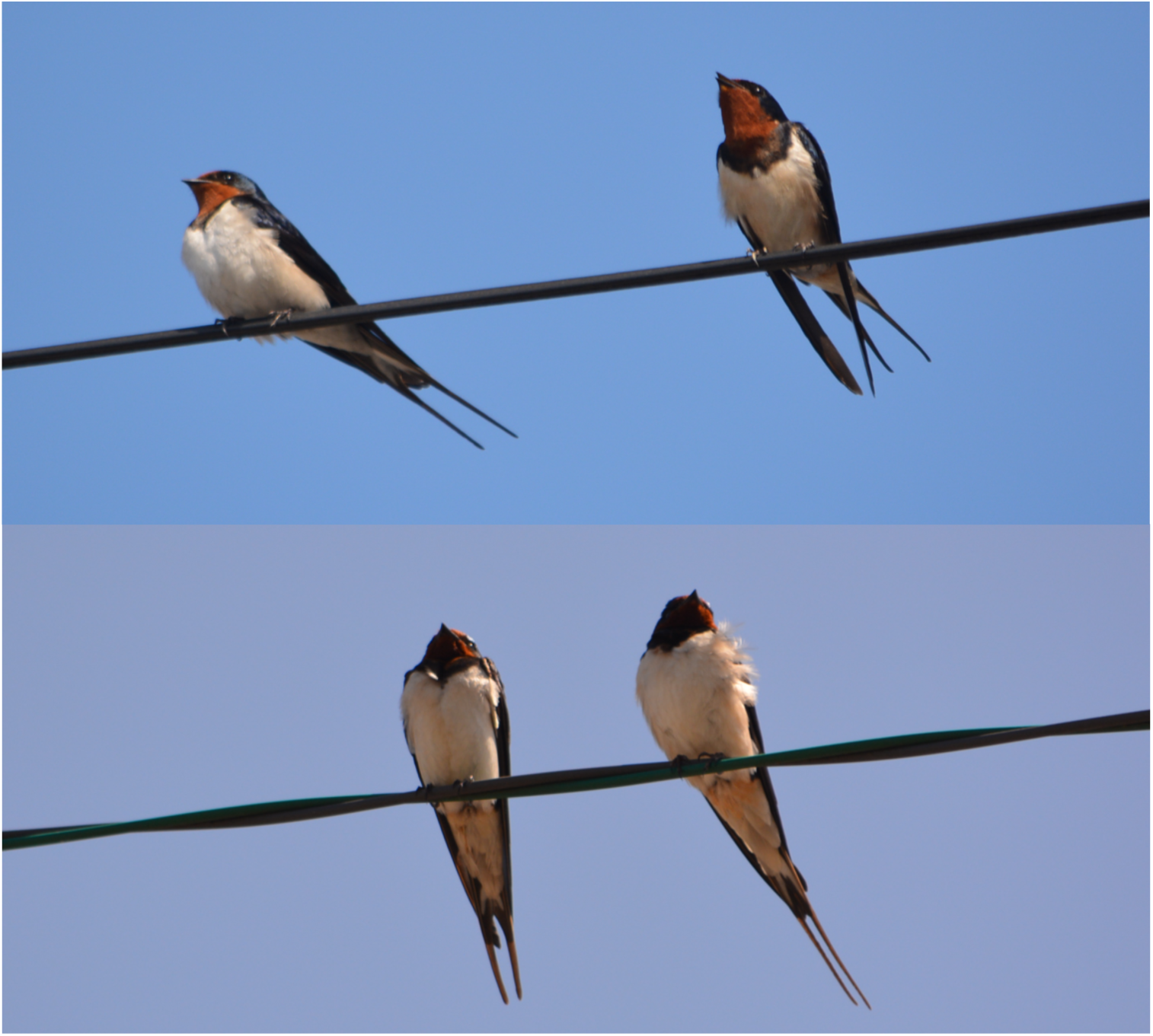
A naturally long-tailed female (upper photograph, left) that resembles the male mate (upper photograph, right; note the color difference in their throats) and a female with a somewhat shorter tail (lower photograph, left) than the male mate (lower photograph, right)

## SUPPLEMENTARY VIDEOS

Video clips showing male copulation attempts and male pairing displays toward stuffed female models: a) male copulation attempt toward a control model; b) male pairing display toward a control model; c) male copulation attempt toward a tail-elongated model; d) male pairing display toward a tail-elongated model.

**Table S2.**
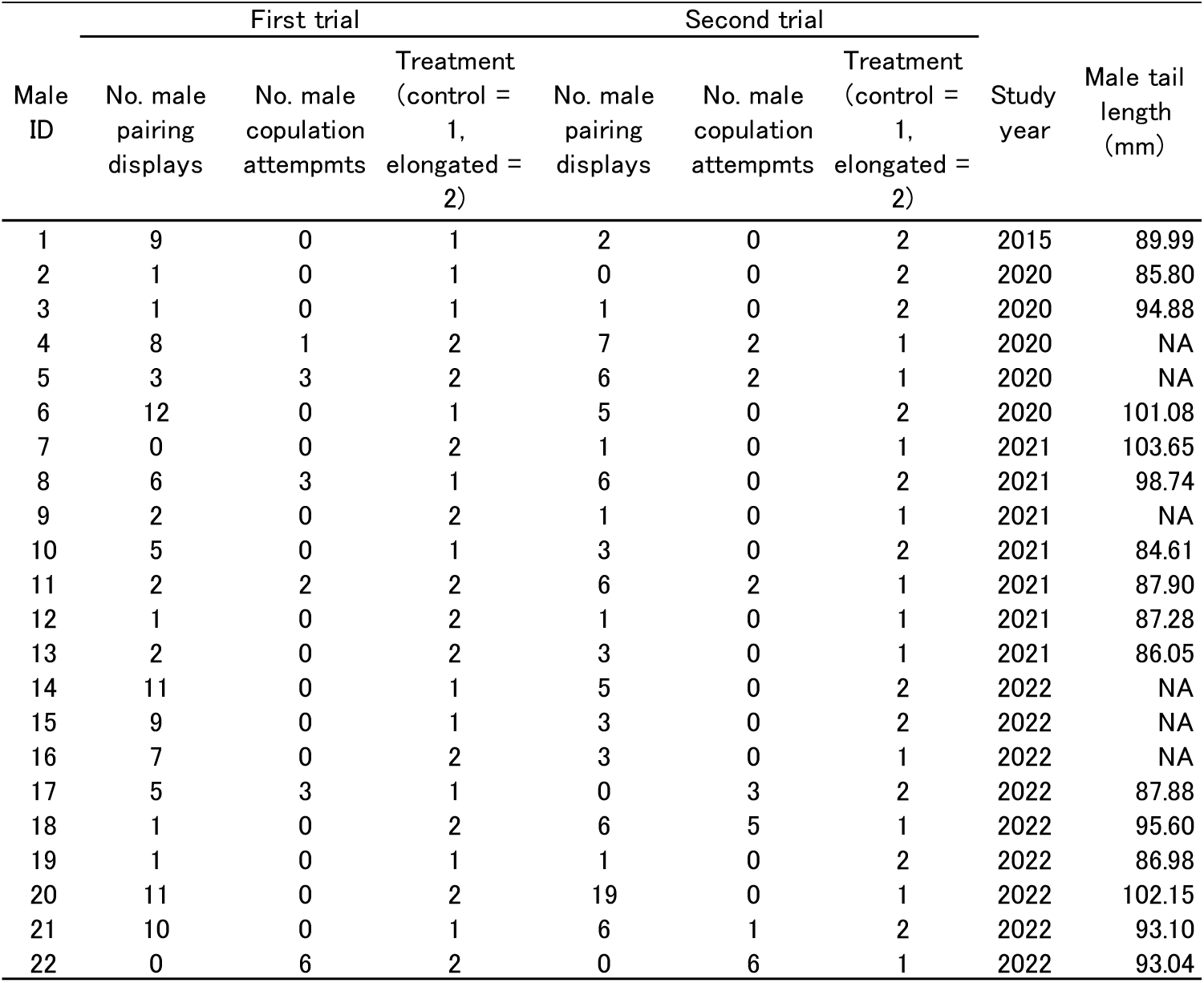
Dataset of the current study

